# Observation-masked neural posterior estimation for heterogeneous epidemiological surveillance data

**DOI:** 10.64898/2026.08.03.742452

**Authors:** Renata Retkute, Christopher A. Gilligan

## Abstract

Disease surveillance data are often sparse, irregularly timed and heterogeneous across observational units, creating challenges for inference in mechanistic epidemiological models. We present observation-masked neural posterior estimation (OM-NPE), a simulation-based Bayesian inference framework for such settings. The approach represents observations on a common temporal grid and records observation availability through a binary mask, enabling heterogeneous surveillance records to be analysed using a single amortised neural posterior estimator. We demonstrate OM-NPE using two contrasting epidemiological systems. First, we fit an effective SEIR model of Zika virus transmission to weekly sentinel surveillance data from six French Polynesian archipelagos during the 2013–14 outbreak. Second, we fit a stochastic compartmental model of *Xylella fastidiosa* spread to annual disease-severity observations from 17 olive groves in Apulia, Italy, each surveyed only two or three times over seven years. In both applications, OM-NPE recovered epidemiological parameters consistent with the assumed epidemiological and observation models, appropriately represented uncertainty, and generated posterior distributions in well under one second without retraining or Markov chain Monte Carlo sampling. Observation masking extends amortised Bayesian inference to heterogeneous longitudinal surveillance systems and supports rapid evaluation of alternative monitoring schedules.

## 1 Introduction

Disease surveillance is central to understanding and managing epidemic and endemic diseases. Mechanistic models provide a formal framework for linking surveillance data to the underlying biological processes governing pathogen transmission, disease progression and recovery, enabling estimation of unobserved epidemiological parameters and supporting prediction and intervention design [14, 1, 11]. In practice, however, surveillance programmes rarely follow ideal sampling schemes; observations are often collected at irregular intervals, vary in duration across locations, and provide only partial views of the epidemic trajectory.

Such temporal misalignment arises across a wide range of epidemiological systems. In some cases, reporting systems in different locations become informative at different stages of an outbreak as infection spreads geographically, resulting in staggered observation windows. In others, sampling consists of only a handful of censuses conducted at irregular intervals over many years, leaving large temporal gaps [3, 4, 15, 27, 24]. Although biologically distinct, these scenarios generate the same fundamental inferential problem: available records provide incomplete temporal coverage that varies systematically between observational units.

Bayesian inference for mechanistic epidemic models has traditionally relied on computationally demanding methods such as data-augmentation Markov chain Monte Carlo, particle filtering and approximate Bayesian computation (ABC), particularly when likelihood functions are unavailable or expensive to evaluate [7, 2, 16, 19]. Hybrid frameworks such as ABC-RF-rejection [23] have improved efficiency by focusing simulations on promising regions of parameter space, yet inference typically remains tied to repeated simulation for each new dataset—a bottleneck that limits practical applicability.

Recent advances in simulation-based inference (SBI) offer an alternative strategy for Bayesian computation in models where simulation is feasible but likelihood evaluation is difficult [6]. Rather than approximating the likelihood indirectly, SBI methods learn relationships between parameters and simulated observations. Among these, neural posterior estimation (NPE) trains a conditional density estimator to approximate the posterior directly from parameter–data pairs generated under the model [20, 12]. Once trained, the resulting estimator is amortised: it can be applied to new observations without further simulation, optimisation or MCMC sampling. Recent work has demonstrated NPE for stochastic epidemic models with final outcomes [17], compartmental and phylodynamic frameworks [22], and viral sequence data [10]. A common thread across these applications, however, is that observations must be representable in a fixed-dimensional format — an assumption that fails for the irregularly timed, location-specific records that characterise many real-world data-collection efforts.

We demonstrate OM-NPE on two contrasting case studies that exemplify the challenges of irregular data. The first uses weekly sentinel returns from six French Polynesian archipelagos during the 2013–14 Zika outbreak, where reporting began at different stages of the epidemic wave. The second uses annual disease-severity assessments from 17 *Xylella fastidiosa*-infected olive groves in Apulia, Italy, where each grove was surveyed only two or three times over a seven-year period. Together, these systems operate on very different temporal and spatial scales, providing complementary tests of the framework’s flexibility. We show that observation masking extends neural posterior estimation to sparse and heterogeneous observational settings while preserving the sub-second posterior evaluation and computational efficiency that make simulation-based inference attractive. The trained estimator also supports rapid exploration of alternative observation schedules without retraining—a capability that would otherwise require repeated model fitting and that has direct practical value for designing efficient sampling protocols.

## 2 Methods

### 2.1 Overview

Our objective is to perform Bayesian inference for mechanistic epidemiological models from longitudinal surveillance data collected under heterogeneous observation schedules. Let *θ* denote the vector of model parameters and let *y* denote observations collected through a surveillance programme. Bayesian inference seeks the posterior distribution

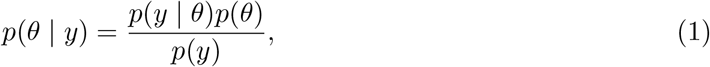

where *p*(*θ*) is the prior distribution and *p*(*y* | *θ*) is the likelihood. For many epidemiological models the likelihood is unavailable in closed form or computationally expensive to evaluate. We therefore adopt a simulation-based inference framework. Following previous applications of neural posterior estimation (NPE) to epidemic models [20, 12, 22, 17], we approximate the posterior directly using a neural density estimator trained on simulated parameter–data pairs generated from the model. The methodological contribution of this paper is not the inference engine itself, but a representation of irregular surveillance data that allows a single amortised posterior estimator to be applied across observational units with different observation schedules. We refer to this framework as observation-masked neural posterior estimation (OM-NPE).

### 2.2 Observation-masked representation

A central challenge is that observational units are often monitored at different times and for different durations. Standard neural density estimators require fixed-dimensional inputs and therefore cannot directly accommodate heterogeneous observation schedules. We address this by representing each surveillance record on a common temporal grid. Let

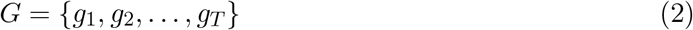

denote a predefined set of candidate observation times. For a surveillance record observed at only a subset of these times we construct two vectors:

1. an observation vector, containing observed values on the grid;
2. a binary observation mask indicating whether each grid point was observed. The resulting input is

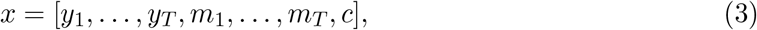

where *m*_*t*_ = 1 if the corresponding observation is available and *m*_*t*_ = 0 otherwise, and *c* denotes any contextual variables associated with the observational unit. The mask distinguishes unobserved time points from genuinely observed zero values. This distinction is critical because zero-filling alone does not allow the network to determine whether a value was observed or absent. Context variables can also be included within the same framework. Throughout this study we include population size as an additional covariate, allowing a single trained network to condition posterior uncertainty on the scale of the observational unit. The observation mask forms part of the network input during both training and inference. The procedure used to generate training examples with heterogeneous observation schedules is described below for each case study.

### 2.3 Neural posterior estimation

Training data are generated by repeatedly sampling parameter values from the prior and simulating observations from the mechanistic model:

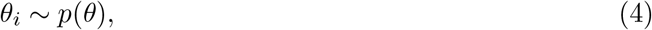

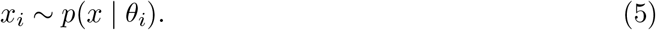

The collection of simulated pairs

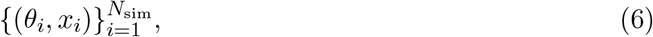

is used to train a conditional density estimator

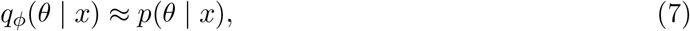

where *ϕ* denotes neural-network parameters. The model is trained by minimising the negative log posterior density

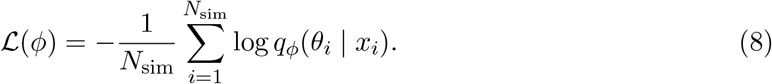

Training is performed only once. Thereafter, the posterior for any observed dataset is obtained by conditioning the trained flow on the corresponding input vector

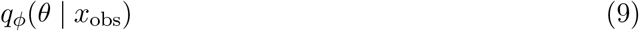

without further simulation or optimisation.

### 2.4 Implementation

We use a Masked Autoregressive Flow (MAF; [21]) as the conditional density estimator, trained via SNPE-C [12] using the Python `sbi` library [26]. A BoxUniform proposal is placed over ***θ*** with bounds chosen to contain the prior’s effective support.

### 2.5 Posterior calibration assessment

To assess the accuracy of the posterior estimator, we employed simulation-based calibration (SBC) [5, 25] to assess performance of individual parameters and tests of accuracy with random points (TARP) [18] to assess multi-parameter performance by the OM-NPE estimation framework. For both diagnostics, we generated *K* = 1,000 calibration samples (***θ***_*k*_, **x**_*k*_) from the prior predictive distribution, and for each drew *L* = 1,000 posterior samples from the trained estimator 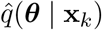.

For SBC, we computed the rank of the true parameter ***θ***_*k*_ among the *L* posterior draws. Under a well-calibrated posterior, these ranks are uniformly distributed on *{*0, …, *L}*; we assessed this by comparing the empirical rank cumulative distribution function (CDF) against the 95% analytic Beta confidence envelope [25]. For TARP, we evaluated expected coverage by sampling reference points from the prior and computing, for each calibration sample and reference point, the fraction of posterior samples that were closer to the reference point than the true parameter [18]. Under calibration, these fractions are uniformly distributed across credibility levels. We summarised global calibration via the area-to-curve (ATC), where positive values indicate overdispersion and negative values indicate underdispersion, and the Kolmogorov–Smirnov (KS) test for uniformity, with *p >* 0.05 indicating no evidence of miscalibration.

### 2.6 Case study I: Zika virus in French Polynesia

#### 2.6.1 Data

The first case study considers the 2013–14 Zika epidemic in French Polynesia. Surveillance data consisted of weekly counts of Zika-like illness reported by sentinel general practitioners (GPs) across six archipelagos over a 25-week window spanning 11 October 2013 to 28 March 2014 [3, 4, 15]. The sentinel network comprised 162 GP practices in total, distributed across archipelagos as follows: 99 in Tahiti, 25 in the Îles-sous-le-Vent, 12 in Mooréa, 13 in the Tuamotu–Gambier archipelago, 8 in the Marquises, and 5 in the Australes. In any given week, only a subset of practices submitted a report; the fraction of active sites varied from week to week within each archipelago. A week was considered unobserved (mask *m*_*t*_ = 0) when no sentinel practice submitted a return. Table 1 summarises the missingness structure by archipelago. The heterogeneity in surveillance coverage across archipelagos, in terms of network size, active reporting fraction, epidemic timing, and missing weeks, makes this dataset a natural test case for the OM-NPE framework.

**Table 1.**
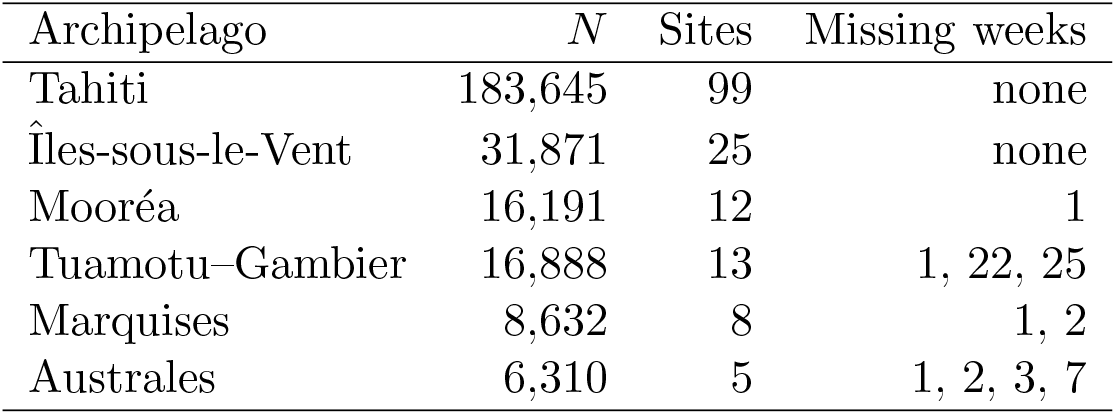
Sentinel surveillance summary by archipelago. Here *N* : resident population; sites: total sentinel GP practices; missing weeks: weeks with no sentinel coverage (*m*_*t*_ = 0).

#### 2.6.2 Transmission model

We implemented the effective SEIR model proposed by Kucharski *et al*. [15], and Funk *et al*. [8] adopting the same compartmental structure and transmission dynamics as the original implementation in [9]. The model incorporates both human and vector dynamics through an effective latent period and is defined by

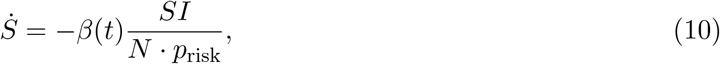

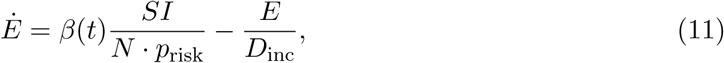

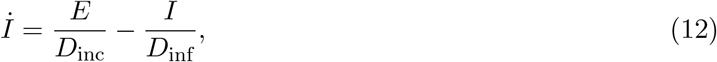

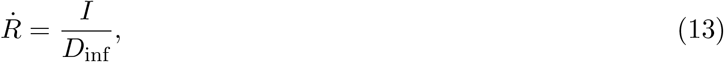

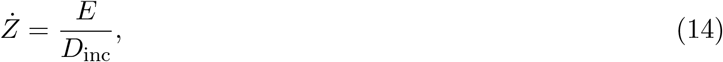

where *D*_inc_ and *D*_inf_ denote the intrinsic incubation and infectious periods respectively, in weeks; *p*_risk_ is the fraction of the population at risk; and *Z* is the cumulative incidence accumulator, which is reset at weekly intervals to obtain weekly case counts. The time-varying transmission rate is given by

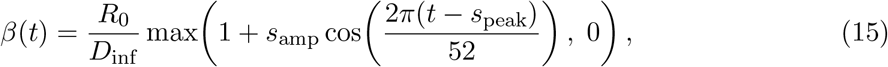

where *R*_0_ is the basic reproduction number, *s*_amp_ controls the amplitude of seasonal forcing, and *s*_peak_ is the week of peak transmission. This seasonal formulation reflects the austral-summer peak in *Aedes* mosquito abundance in French Polynesia.

Initial conditions are set as

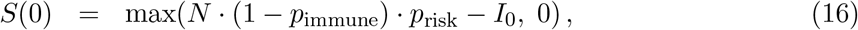

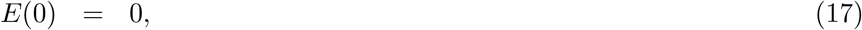

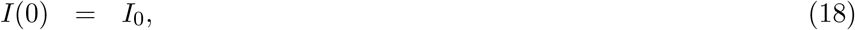

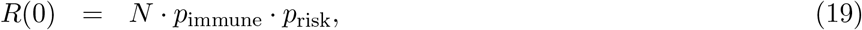

where *p*_immune_ is the initial seroprevalence and *I*_0_ is the number of initially infectious individuals at the start of the simulated epidemic. To account for uncertain epidemic onset timing across archipelagos, we introduce an offset parameter *t*_0_, sampled uniformly from the prior [0, 10] weeks. The observation window is treated as a fixed *T*_max_ = 25-week slice beginning at time *t*_0_ within a longer simulated epidemic of total duration *T*_total_ = *T*_max_+*T*_0,max_ = 35 weeks, where *T*_0,max_ = 10.

The reporting process is modelled as

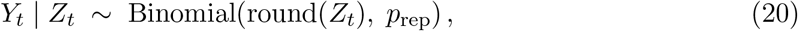

where *Z*_*t*_ is the number of new infectious cases in week *t* and *p*_rep_ is the reporting probability. This replaces the truncated-Gaussian observation model used in the original MCMC implementation [15]; the exact Binomial is appropriate here because the neural network is trained on simulated data and does not require the computational expediency of a continuous approximation. This is particularly important for the tails of the epidemic, where small case counts make the Gaussian approximation unreliable.

The inferred parameter vector is therefore

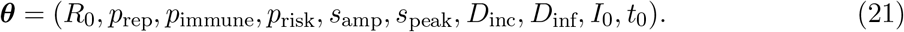

Parameter ranges are given in SI Table SI1.

### 2.7 Case study II: *Xylella fastidiosa*

#### 2.7.1 Data

The second case study considers disease monitoring data from 17 olive (*Olea europaea*) groves infected with *Xylella fastidiosa* subsp. *pauca* in Puglia, southern Italy [27]. The dataset comprises visual assessments of disease severity for each tree, scored on a 0–5 canopy desiccation scale. Scores were classified into three epidemiological compartments: score 0 as susceptible or symptomless infected (*S* +*I*_*A*_), scores 1–3 as symptomatic infected (*I*_*S*_), and scores 4–5 (majority or all branches dead) as desiccated (*I*_*D*_).

The data came from 2,959 olive trees across 16 infected groves (mean plot area 1.86 *±* 1.57 ha; mean tree density 97.9 *±* 41.7 trees/ha), each surveyed at two or three time points between 2016 and 2018 [28, 13]. An additional grove of 187 trees was surveyed at an earlier stage of infection in January 2014 and April 2015 [27], prior to the main survey campaign. Table 2 summarises the 17 plots by identifier, grove size, and survey years.

**Table 2.** Olive grove plots used for *Xylella fastidiosa* model fitting [27] *N* : number of trees surveyed.

| Plot | $N$ | Survey years |
| --- | --- | --- |
| A20 | 521 | 2016, 2017, 2018 |
| C20 | 441 | 2016, 2017, 2018 |
| B51 | 335 | 2016, 2017 |
| B11 | 219 | 2016, 2017 |
| C2 | 190 | 2016, 2017, 2018 |
| A4 | 185 | 2016, 2017 |
| B3 | 160 | 2016, 2017, 2018 |
| C22 | 136 | 2016, 2017 |
| B8 | 131 | 2016, 2017, 2018 |
| B1 | 98 | 2016, 2017 |
| A9 | 89 | 2016, 2017 |
| B20 | 82 | 2016, 2017 |
| B22 | 60 | 2016, 2017 |
| B21 | 54 | 2016, 2017, 2018 |
| A5 | 40 | 2016, 2017 |
| B200 | 31 | 2016, 2017 |
| MC | 187 | 2014, 2015 |
| Total | 2,959 |  |

Because the initial infection date of each grove was unknown, the time of infection was treated as a latent variable during model fitting, with simulations run for up to 12 years — the estimated upper bound on the duration of infection in the demarcated zone [27]. Within this 12-year window, each plot contributes observations at only 2–3 irregularly spaced time points, leaving the remainder of the window unobserved. Survey timing also differed between plots, with some visited in June and others in October or December within the same calendar year. The combination of unknown infection onset, sparse temporal coverage, and plot-specific visit schedules makes this dataset an extreme case of heterogeneous longitudinal surveillance, and a natural test of the OM-NPE framework’s ability to pool information across observational units with disparate observation schedules.

#### 2.7.2 Transmission model

We use the discrete-time compartmental model of White *et al*. [27]. Each olive tree can be in one of four states: susceptible (*S*), asymptomatic infected (*I*_*A*_), symptomatic infected (*I*_*S*_), and desiccated (*I*_*D*_), with annual transitions:

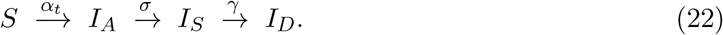

The annual infection probability is

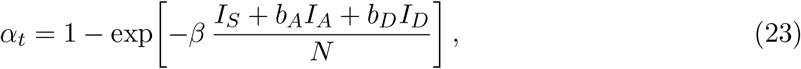

where *β* is the effective contact rate (expected number of *Xylella*-transmitting contacts per year from each symptomatic tree), and *b*_*A*_, *b*_*D*_ *∈* [0, 1] are the infectivities of asymptomatic and desiccated trees relative to symptomatic trees, respectively. Asymptomatic trees are expected to have low infectivity owing to reduced bacterial load, whilst desiccated trees may be less attractive to the insect vector [27]. The symptom-appearance probability 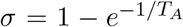 and desiccation probability 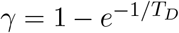 are derived from exponential waiting times, where *T*_*A*_ is the mean duration of the asymptomatic period (years) and *T*_*D*_ is the mean time to desiccation after the minimum symptomatic period has elapsed (years).

Additionally, to ensure biologically realistic disease progression [27], a desiccation-delay parameter *τ* enforces a minimum sojourn of *τ* years in *I*_*S*_ before trees can enter *I*_*D*_, capturing the observed lag between early symptom expression and rapid branch dieback; following [27] this is fixed at *τ* = 3 years. The initial proportion of asymptomatic infected trees *I*_*A*,0_ is treated as a free parameter because the calendar date of infection for each grove is unknown. The inferred parameter vector is therefore

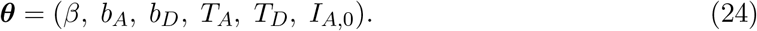

Parameter ranges are given in SI Table SI2

## 3 Results

### 3.1 Inference from heterogeneous surveillance records: Zika virus in French Polynesia

We first evaluated the performance of the trained amortised NPE on the French Polynesia outbreak data. To ensure the posterior generalises to arbitrary observation schedules, we trained the neural network using a simulation-based masking strategy. For each training example, the number of observed weeks *n*_obs_ was drawn uniformly from *{*1, …, 25*}*, and the observed positions were selected uniformly at random without replacement from the 25-week window.

This exposes the network to the full space of missingness patterns, enabling amortised inference for any surveillance schedule encountered at test time, including the irregular, archipelago-specific reporting patterns present in the French Polynesia data.

We trained a single-round NPE using 10^6^ simulations. For each simulation, parameters ***θ*** were drawn from the priors described above; a population size *N* was sampled uniformly from *{*1,000, …, 250,000*}*; and the simulated weekly incidence was divided by *N* to yield per-capita observations. The input to the network is a 51-dimensional vector

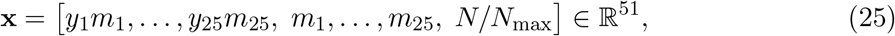

where unobserved entries are set to zero by the binary observation mask, the mask itself is appended to allow the network to distinguish a true zero from a missing observation, and the scaled population *N/N*_max_ with *N*_max_ = 250,000 is provided as a scalar context feature. Training required approximately 87 minutes on a single CPU.

Posterior distributions were obtained for all six archipelagos using the same trained estimator, conditioned on each region’s unique mask and observed case series; for each region, we drew 1,000 samples from the posterior, a process that required approximately 0.3 seconds in total. Figure 1 compares the observed weekly case counts with the posterior predictive distributions for each archipelago, showing that the model captures the key features of the epidemic trajectories across all regions.

**Figure 1.**
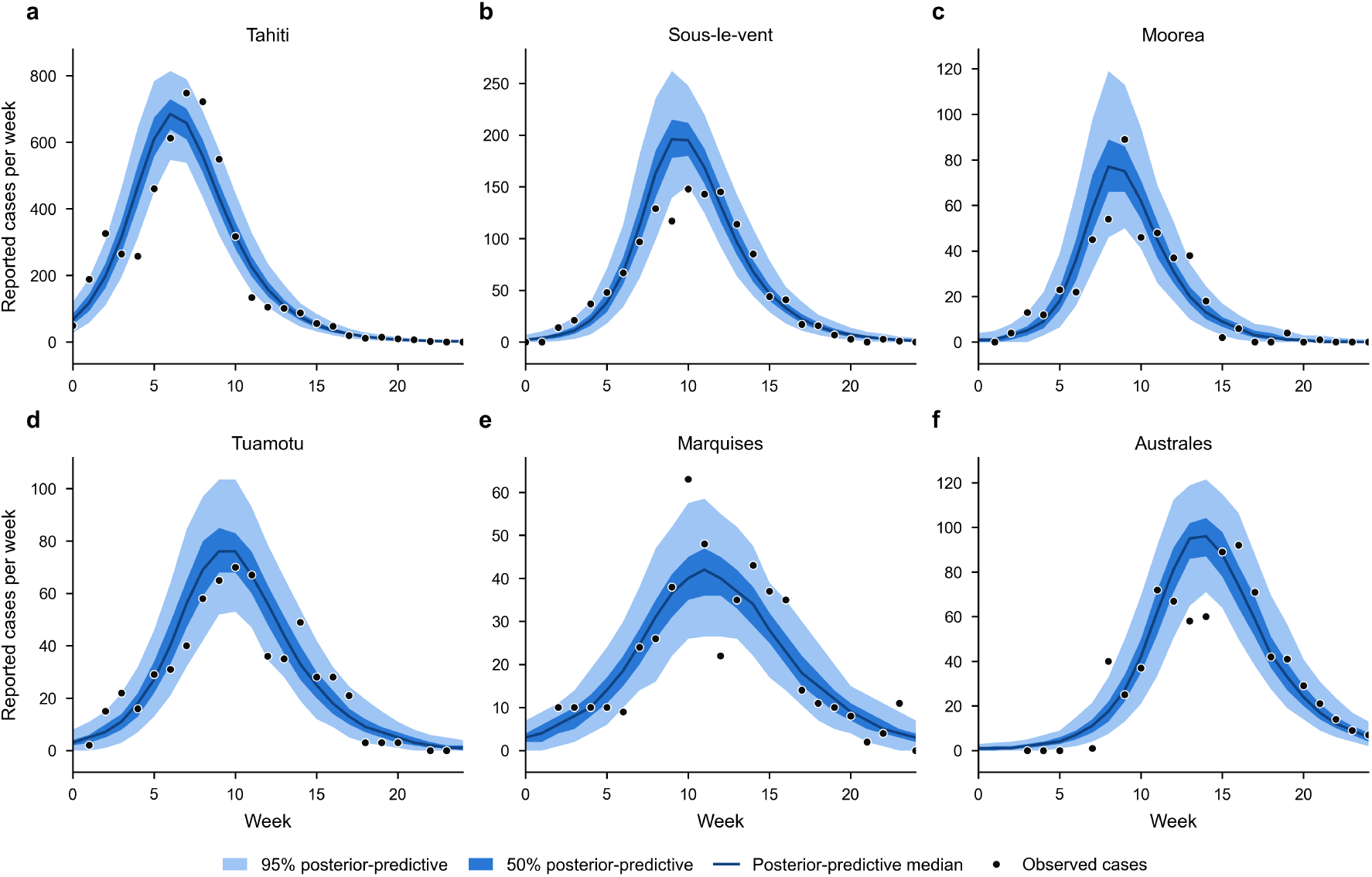
Comparison of reported cases and fitted model trajectories. Blue dots show weekly reported Zika virus cases [15]; dark blue line shows the median of 1,000 posterior samples from the fitted NPE; the darker shaded region shows the 50% credible interval; and the lighter shaded region shows the 95% credible interval.

Simulation-based calibration over *K* = 1,000 prior draws confirmed well-calibrated posteriors across all ten model parameters, with rank CDFs falling within the 95% confidence band of the uniform null distribution (SI Fig. 2A). The absence of systematic S-shaped or bowed departures indicates that the amortised estimator produces posteriors with neither appreciable overconfidence nor directional bias under the reporting schedules observed across Pacific island territories. The TARP KS-test p-value of 1.00 indicates that the distribution of TARP fractions is consistent with uniformity, confirming that the posterior estimator does not exhibit structural miscalibration. The area-to-curve of 0.28 reflects mild, homogeneous overdispersion—the posterior is systematically conservative—which is preferable to underconfidence as it avoids overstating parameter certainty ( SI Fig. 2B).

As an additional check, we compared our *R*_0_ estimates to those reported in Kucharski *et al*.[15], who analysed the same data using MCMC. The estimates of the basic reproduction number were broadly consistent across archipelagos, with posterior medians ranging from approximately 2.4 to 4.7 (Table 3). Median values and 95% credible intervals were very close to the estimated values in [15]. The closest agreement was observed for Tahiti, Moorea, and Tuamotu–Gambier, for which the difference between our median *R*_0_ estimate and the MCMC median reported in [15] was within 0.1. The largest difference was for the Australes, where our median was 0.4 lower (2.7 vs. 3.1) and the 95% credible interval was slightly narrower (1.8–4.0 vs. 2.2–4.6), though the estimates remained in broad consensus. Supplementary Figure 1 shows the corresponding posterior estimates for all model parameters across the six archipelagos.

**Table 3.** Estimates for the basic reproduction number, *R*_0_ using NPE (this study) and in Kucharski *et al*. [15].

| Archipelago | This study | Kucharski et al. 2016 |
| --- | --- | --- |
| Tahiti | 3.5 (2.4–5.0) | 3.5 (2.6–5.3) |
| Îles-sous-le-Vent | 3.7 (2.1–6.6) | 4.1 (3.1–5.7) |
| Mooréa | 4.7 (2.4–9.8) | 4.8 (3.2–8.4) |
| Tuamotu–Gambier | 3.1 (2.0–5.4) | 3.0 (2.2–6.1) |
| Marquises | 2.4 (1.7–3.6) | 2.6 (1.7–5.3) |
| Australes | 2.7 (1.8–4.0) | 3.1 (2.2–4.6) |

### 3.2 Sequential inference

A key operational advantage of the amortised OM-NPE framework is that posterior inference requires no retraining or additional simulation once the estimator is trained. To demonstrate this, we conducted a retrospective sequential inference experiment across all six French Polynesian archipelagos. Starting from week 3, we progressively revealed one additional week of surveillance data at a time and re-ran the trained posterior estimator at each step, tracking the evolution of posteriors for *R*_0_, reporting probability *p*_rep_, and epidemic onset delay *t*_0_. Each inference call took under one second, so the full sequential analysis for all archipelagos was completed in minutes.

Posteriors for *R*_0_ converged rapidly once the ascending phase of the epidemic was captured in the observation window. For Tahiti, the 90% credible interval for *R*_0_ excluded 1 after approximately seven weeks of data — coinciding with the epidemic peak — providing a statistically grounded declaration of sustained transmission well before the epidemic had resolved. Smaller archipelagos (Marquises, Australes) required proportionally more of the observation window to achieve the same threshold, reflecting the lower signal-to-noise ratio in smaller populations. Posterior uncertainty in *p*_*rep*_ and *t*_0_ remained elevated until post-peak data were available, consistent with the identifiability of these parameters depending on the full epidemic trajectory rather than early ascending-phase counts alone.

### 3.3 Inference from sparse surveillance: *Xylella fastidiosa* surveys

We next applied OM-NPE to disease-severity observations from 17 *X. fastidiosa* infected olive groves in Apulia. For each grove, annual disease-severity observations were mapped onto an annual grid spanning years 1 - 10 after epidemic establishment. Each observation consisted of the observed proportions

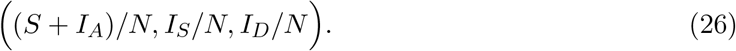

Years without surveillance were represented through the observation mask. Population size was included as a context variable so that posterior uncertainty reflected finite-population stochasticity across groves of different sizes.

We trained a single-round NPE using 10^6^ simulations. For each simulation, parameters ***θ*** were drawn from the priors; a population size *N* was sampled uniformly from *{*25,…,600*}* and the simulated annual proportions were divided by *N* to yield per-capita observations. The input to the network is a 29-dimensional vector

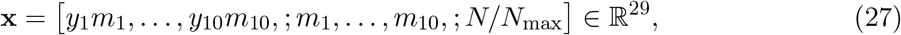

where unobserved entries are set to zero by the mask, the mask itself is appended to allow the network to distinguish a true zero from a missing observation, and the scaled population is provided as a scalar context feature. Training examples were constructed from a mixture of two observation-schedule distributions. With probability 0.5, the observation mask matched one of the survey schedules observed in the 17 olive groves. With probability 0.5, the mask was generated by selecting between two and seven observation years from the 7-year grid at random. The empirical schedules ensured good coverage of the surveillance histories represented in the study dataset, whereas the random schedules exposed the estimator to a wider range of temporal observation patterns. Consequently, the trained estimator could be used both for inference on the observed groves and for evaluating alternative surveillance schedules. Training required approximately 45 minutes on a single CPU.

Once the NPE was trained, we drew 100 samples from the posterior for each grove. . Despite the sparse observation schedule, the estimator produced informative posterior distributions for all groves (SI Figure 3), with fitted trajectories closely matching the observed surveillance data (Fig.3A). Simulation-based calibration revealed well-calibrated posteriors for *β* and *T*_*D*_, with rank CDFs falling within the 95% confidence band of the uniform null distribution (SI Fig. **??**A). Mild departures were observed for *T*_*A*_ and *IA*_0_, while *b*_*A*_ and *b*_*D*_ showed more pronounced deviations. While the TARP KS-test p-value of 1.00 affirms that the estimator adheres to the correct distributional shape, the negative area-to-curve (-0.21) signals a subtle, uniform tendency toward underdispersion, implying that the posterior is marginally overconfident but not structurally mis-calibrated ( SI Fig. 4B).

**Figure 2.**
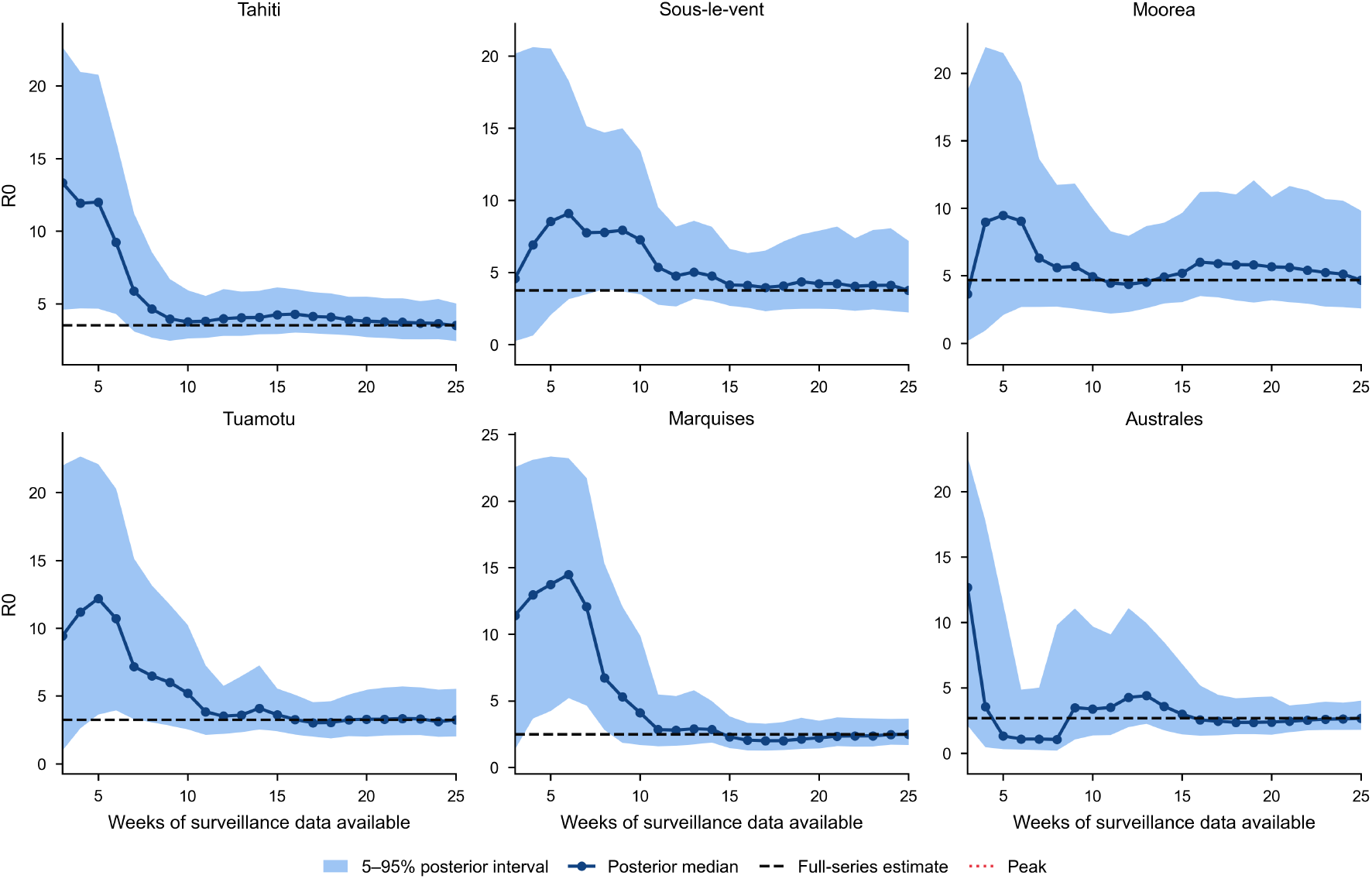
Sequential posterior inference for the basic reproduction number *R*_0_ for Zika virus across six French Polynesian archipelagos. Lines show the posterior median and shaded bands the 90% credible interval as the number of available weekly surveillance observations increases from 3 to 25 weeks. Dashed horizontal lines marks median estimate based on all data.

**Figure 3.**
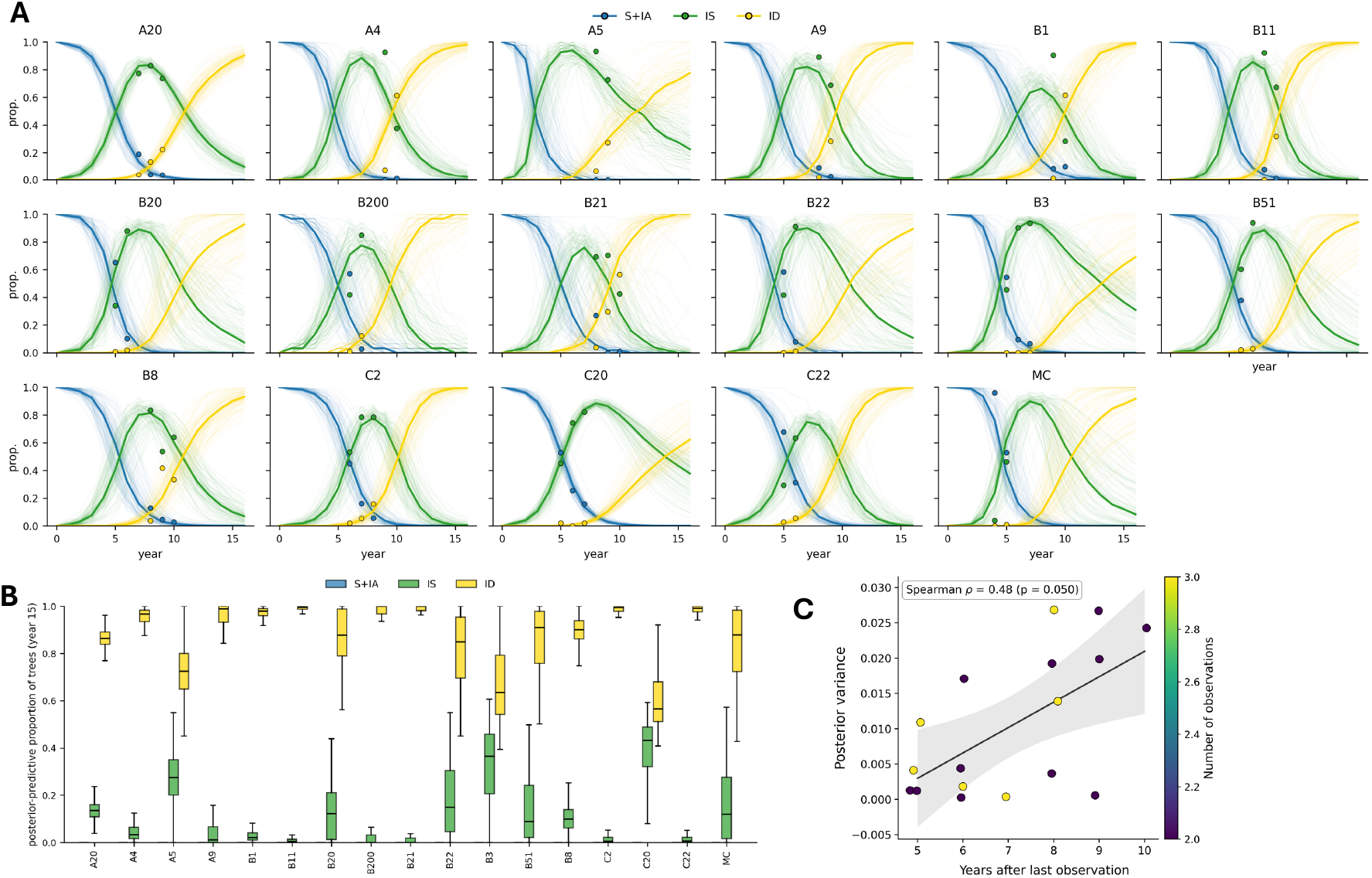
(A) Comparison of reported cases and fitted model trajectories using the OM-NPE framework for *X. fastidiosa* groves in Puglia. Dots show surveillance data [27], lighter lines show individual posterior trajectories, and darker line shows median. (B) Predicted proportion of healthy + asymptomatic, symptomatic, and desiccated trees 16 years from infection estimated using the OM-NPE framework. (C) Posterior forecast uncertainty for the proportion of desiccated trees versus extrapolation horizon, across all monitored plots. The solid line and shaded band show the ordinary-least-squares fit and its 95% confidence interval

To examine the long-term epidemic trajectory, we evaluated the fitted model at year 15 post-infection (Fig.3B). The estimated proportions revealed marked inter-grove heterogeneity, reflecting differences in inferred epidemiological parameters and observation histories. These inferences are conditional on the assumed observation model, which treats disease-severity classifications as error-free and does not explicitly account for observer mis-classification or differences in within-year survey timing. A notable multi-modal pattern emerged across groves. Groves A20, A4, A9, B1, B11, B200, B21, C2, and C22 exhibited high desiccated proportions (median 0.8–0.95) with narrow credible intervals, indicating early infection and rapid progression to terminal desiccation. In contrast, groves A5, B20, B22, B23, B3, B51, C20, and MC showed considerably wider posterior spreads and elevated symptomatic fractions, consistent with later infection onset or prolonged residence in the symptomatic phase. This bimodal distribution highlights the critical role of stochastic variation in initial conditions and the timing of disease establishment. To assess if observation gaps influenced forecast reliability, we examined the relationship between posterior-predictive uncertainty and extrapolation distance from the final survey to year 16 (Fig.3C). Larger gaps showed a suggestive trend towards greater forecast uncertainty (Spearman’s *ρ* = 0.48, *p* = 0.05), reflecting the inherent challenge of long-term prediction from sparse surveillance data.

### 3.4 Optimising surveillance schedules

The observed relationship between extrapolation distance and forecast uncertainty raises a practical question for disease managers: given limited monitoring resources, how should observations be scheduled to maximise forecast reliability? To address this, we followed the practice of White *et al*. [27] by aggregating data from the 17 groves into a single synthetic observation by summing the counts of trees in each disease compartment across all years where data were present and calculating the corresponding proportions. This aggregated observation served as a representative “typical” grove, enabling a systematic assessment of how different surveillance schedules affect predictive uncertainty while controlling for variation in underlying epidemic trajectories.

We generated all possible binary masking patterns for years 4 through 10 (2^7^ = 128 schedules), excluding the trivial all-missing case, and assumed a total population of *N* = 500 trees. For each schedule, we drew 100 posterior samples from the trained NPE and computed the posterior-predictive variance of the proportion of desiccated trees at year 16. Figure 4A displays these schedules ordered by decreasing forecast variance, with observation years indicated by filled squares.

**Figure 4.**
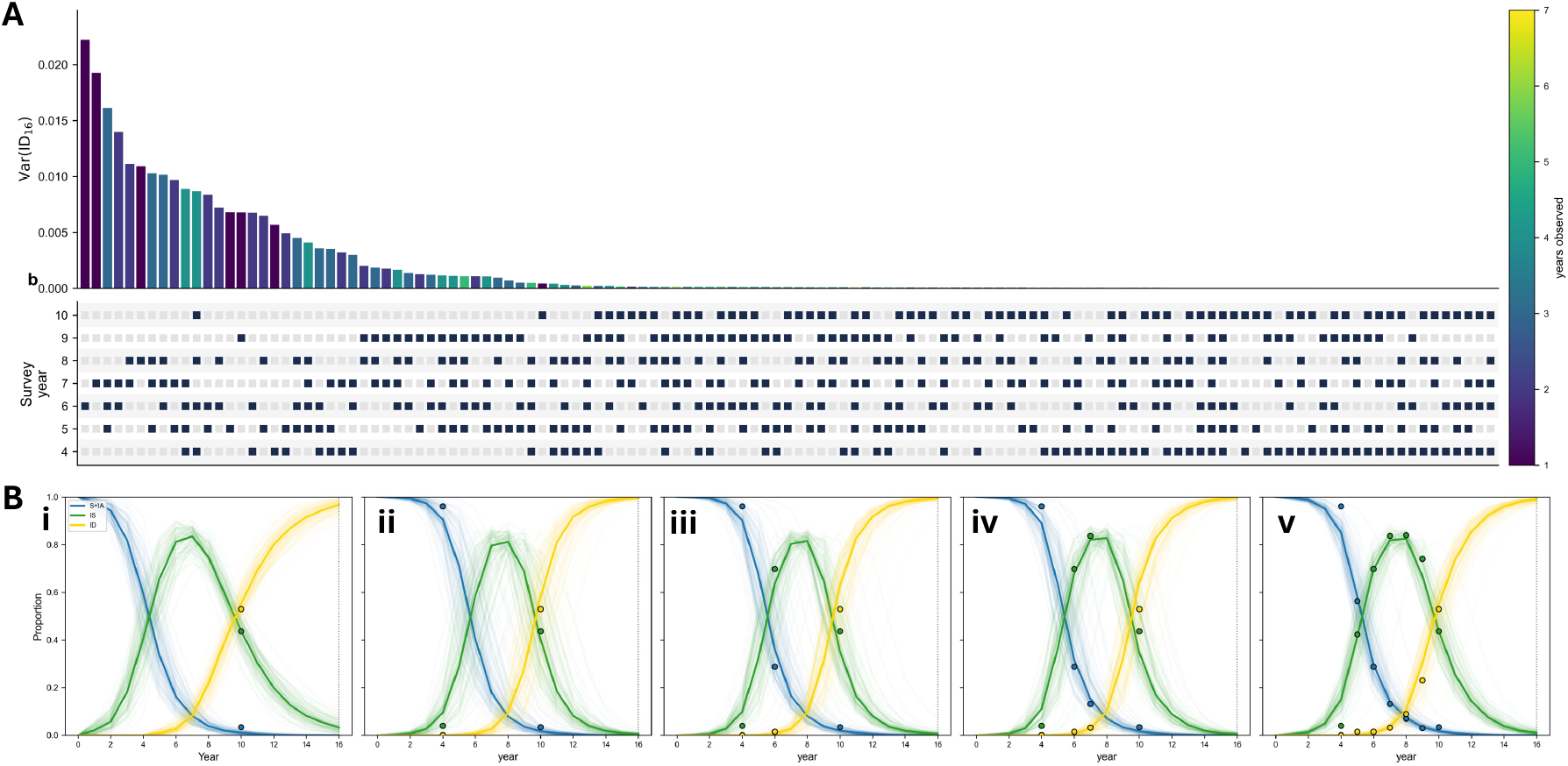
(A) Surveillance schedule optimisation for forecasting *X. fastidiosa* progression. (A) All possible binary masking patterns for years 4-10 (2^7^ = 128 schedules), ordered by decreasing posterior-predictive variance of the proportion of desiccated trees at year 16 (Var(*I*_*D*_(16))). Filled squares indicate observation years; schedules are colour-coded by the number of observations (*n*_obs_). (B) Posterior-predictive fits for the best-performing schedule at each observation count *n*_obs_ = 1, 2, 3, 4 (year 10 (i); years 4, 10 (ii); years 4, 6, 10 (iii); and years 4, 6, 7, 10 (iv)), alongside the fully observed 7-year reference (v).

Forecast variance ranged from 8.70 *×* 10^*−*5^ for the fully observed 7-year schedule to 0.024 for schedules with only a single early observation (year 6), representing a nearly 300-fold difference in predictive uncertainty. For each observed-year count *n*_obs_ = 1, 2, 3, 4 (i.e. the number of years within the 7-year observation window during which disease was observed), we identified the best-performing schedule as the one with the lowest posterior-predictive variance Var(*I*_*D*_(16)) among all schedules of that size, with the fully observed 7-year schedule included as a reference (Fig. 4B). The optimal schedules were year 10 for *n*_obs_ = 1 (variance 3.90 *×* 10^*−*4^), years (4, 10) for *n*_obs_ = 2 (1.95 *×* 10^*−*5^), years (4, 6, 10) for *n*_obs_ = 3 (9.27 *×* 10^*−*6^), and years (4, 6, 7, 10) for *n*_obs_ = 4 (1.35 *×* 10^*−*5^), compared with the fully observed schedule (years 4–10) with variance 8.70 *×* 10^*−*5^. Notably, both the best *n*_obs_ = 3 and *n*_obs_ = 4 schedules yielded lower forecast variance than the fully observed schedule. This result suggests that, for this aggregated example and forecasting objective, strategically timed observations may be more informative than regular annual sampling. The relationship between forecast variance and the number of observations was not strictly monotonic, however, with the best *n*_obs_ = 4 schedule performing slightly worse than the best *n*_obs_ = 3 schedule and both outperforming the fully observed design. This may reflect a combination of Monte Carlo variability arising from the finite number of posterior samples and the fact that observation timing, rather than observation count alone, determines the information available for forecasting.

The systematic evaluation also allowed us to benchmark the NPE against the MCMC-based estimates of *X. fastidiosa* transmission parameters in [27] (Table 4). The NPE posterior medians for *T*_*A*_, *T*_*D*_, and *I*_*A*,0_ were broadly consistent with the MCMC results. However, the transmission rate *β* exhibited a lower median (9.51 vs. 17.88), with comparable credible interval bounds (4.02–30.62 vs. 6.33–24.88). Notably, the NPE estimated a higher infectivity rate for asymptomatic trees (*b*_*A*_ = 0.15 vs. 0.015) and a broadly similar value for desiccated trees (*b*_*D*_ = 0.54 vs. 0.503), though with wider credible intervals in each case. Despite agreement for several parameters, notable differences were observed for *β* and *b*_*A*_. In particular, the lower OM-NPE estimate of *β* was accompanied by a higher estimate of the infectivity of asymptomatic trees (*b*_*A*_; Table 4), consistent with a degree of compensation between these parameters. The broadly similar estimates for the disease-progression parameters *T*_*A*_, *T*_*D*_, and *I*_*A*,0_, however, suggest that the two inference approaches lead to comparable conclusions regarding disease progression while highlighting potential differences in the attribution of transmission.

**Table 4.** Parameter estimates reported as posterior medians with 95% credible intervals from OM-NPE (this study) and compared with the MCMC estimates reported by White *et al*. [27]. *β*: contact rate of symptomatic trees (per year); *b*_*A*_ and *b*_*D*_: infectivity of asymptomatic and desiccated trees relative to symptomatic trees; *T*_*A*_: mean duration of the asymptomatic period (years); *T*_*D*_: mean time to desiccation after the minimum symptomatic period of 3 years has elapsed; *I*_*A*,0_: initial proportion of asymptomatic infected trees.

| Parameter | This study | White et al. (2020) |
| --- | --- | --- |
| $\beta$ | 9.51 (4.02, 30.62) | 17.88(6.33, 24.88) |
| $b_A$ | 0.15 (0.01, 0.64) | 0.015 (0.0, 0.44) |
| $b_D$ | 0.54 (0.08, 0.89) | 0.5 (0.02, 0.98) |
| $T_A$ | 1.35 (1.15, 1.49) | 1.19 (1.09, 1.27) |
| $T_D$ | 1.07 (0.72, 1.41) | 1.36 (1.11, 1.59) |
| $I_{A,0}$ | 0.008 (0.002, 0.015) | 0.0065 (0.003, 0.008) |

## 4 Discussion

The principal contribution of this study is a framework for Bayesian inference from surveillance data collected under heterogeneous observation schedules in different but related regions. Building on recent developments in neural posterior estimation [20, 12, 22, 17], we introduce an observation-mask representation that enables a single amortised posterior estimator to be applied across observational units that differ in the timing, duration and completeness of observation. The approach addresses a practical limitation of simulation-based inference methods, namely the assumption that observations can be represented in a common fixed-dimensional format.

The two case studies demonstrate that the challenge of irregular surveillance arises in very different epidemiological settings. In the French Polynesia Zika epidemic, observation schedules differed because reporting systems became active at different stages of the epidemic wave amongst the archipelagos. In the *X. fastidiosa* system, irregularity arose because different olive groves were observed only a small number of times over a period of several years. Despite these differences in pathogen biology, host system and temporal scale, the same observation-mask framework supported inference in both applications. This suggests that, under the specified priors, observation models and mask-generating distributions, the approach captures a general feature of surveillance data rather than a property of any particular epidemiological model. The extent to which these findings generalise across alternative observation processes and surveillance designs remains an important topic for future work.

The results also illustrate the importance of distinguishing missing observations from observed zeros. In both systems, the observation mask allowed the network to infer whether information was absent because a location was not surveyed or because the observed value was genuinely close to zero. This distinction is often lost when irregular time series are represented through interpolation or simple zero-filling. More generally, the results suggest that explicit representation of observation availability may be as important as the choice of inference algorithm itself when analysing heterogeneous surveillance records.

Recent studies have established neural posterior estimation as a practical tool for mechanistic epidemic models. Kypraios [17] demonstrated accurate amortised inference for stochastic epidemic models observed through final outbreak outcomes, while Pinotti et al. [22] showed that neural posterior estimation can recover posterior distributions for epidemic and phylodynamic models using richer temporal and genetic data. The present work addresses a different problem. Rather than proposing a new inference engine, it extends amortised inference to surveillance datasets in which observation schedules vary amongst observational units. The contribution therefore lies in the representation of the observations rather than in the underlying neural density estimator. By separating the cost of training from the cost of inference, the framework enables near-instantaneous posterior estimation across observational units with heterogeneous surveillance histories.

A practical consequence of the amortised framework is that the same trained estimator can be reused for tasks beyond parameter estimation. In the *X. fastidiosa* application, the estimator was used to explore the consequences of alternative surveillance schedules, to quantify the value of additional observations and to assess the effects of missing data. Such analyses would require repeated model fitting under conventional Bayesian methods but can be performed rapidly once the amortised estimator has been trained. This suggests a broader role for simulation-based inference as a tool for surveillance design as well as parameter estimation.

The two applications also provide insight into the relationship between surveillance intensity and parameter identifiability. In both systems, posterior uncertainty increased predictably as information was removed from the observations. For the Zika case study, uncertainty in epidemic onset increased when early epidemic data were masked. For the *X. fastidiosa* system, parameter recovery depended strongly on the timing of observations. Importantly, posterior uncertainty increased as observational information was removed, indicating that the estimator appropriately reflected differences in data availability across surveillance schedules.

Several methodological directions warrant further investigation. First, the observation schedules used during training were generated by masking a common temporal grid, whereas the empirical surveillance systems analysed here exhibited more structured observation patterns. Future work could examine how different mask-generating distributions affect performance and quantify the robustness of the framework to distribution shifts between training and deployment observation schedules. Second, the observation-mask representation could be compared systematically with alternative approaches for handling irregular longitudinal observations in order to better understand the circumstances under which explicit representation of observation availability provides the greatest benefit. Third, although the calibration results presented here indicate generally well-calibrated posteriors (SI Figs. 2 and 4), with only modest departures for some weakly identified parameters in the sparse *Xylella* case study, further work is needed to understand how calibration is affected by increasing sparsity, structured missingness, and more complex surveillance designs.

Both epidemiological models were deliberately simplified representations of their respective systems. The Zika model treated archipelagos independently and did not explicitly represent inter-island transmission, while the *X. fastidiosa* transmission model considered within-grove dynamics only. Although these simplifications were sufficient to demonstrate the observation-masking framework, future applications may require spatial, hierarchical or network-based models together with richer representations of observations and covariates.

More broadly, the observation-mask framework is not specific to human and plant disease or more generally to infectious disease epidemiology. Any surveillance system that combines mechanistic models with irregular longitudinal observations may face similar challenges. Potential applications include ecological monitoring, wildlife disease surveillance, environmental observation programmes and public-health surveillance systems in which reporting schedules differ across locations. Extending the framework to these settings, and exploring alternative representations for more complex observation structures, are promising directions for future work. An important next step will be to evaluate observation-masked inference in settings with structured missingness patterns, richer covariate information and more complex epidemiological models.

In summary, observation-masked neural posterior estimation extends amortised Bayesian inference to the sparse, irregular and heterogeneous observation schedules that characterise many real-world surveillance systems. The framework retains the computational advantages of simulation-based inference while accommodating variability in observation schedules, providing a practical route to Bayesian inference from imperfect longitudinal data.

## Declarations

### Ethics

This study used only aggregate disease-survey records and required no ethical approval.

### Data availability

The surveillance data, model code, and inference code are available at Github.

### Author contributions

R.R. and C.A.G. designed the study; R.R. implemented the models and inference and analysed the results; R.R. and C.A.G. wrote the manuscript.

## Supplementary Information

**SI Fig. 1:**
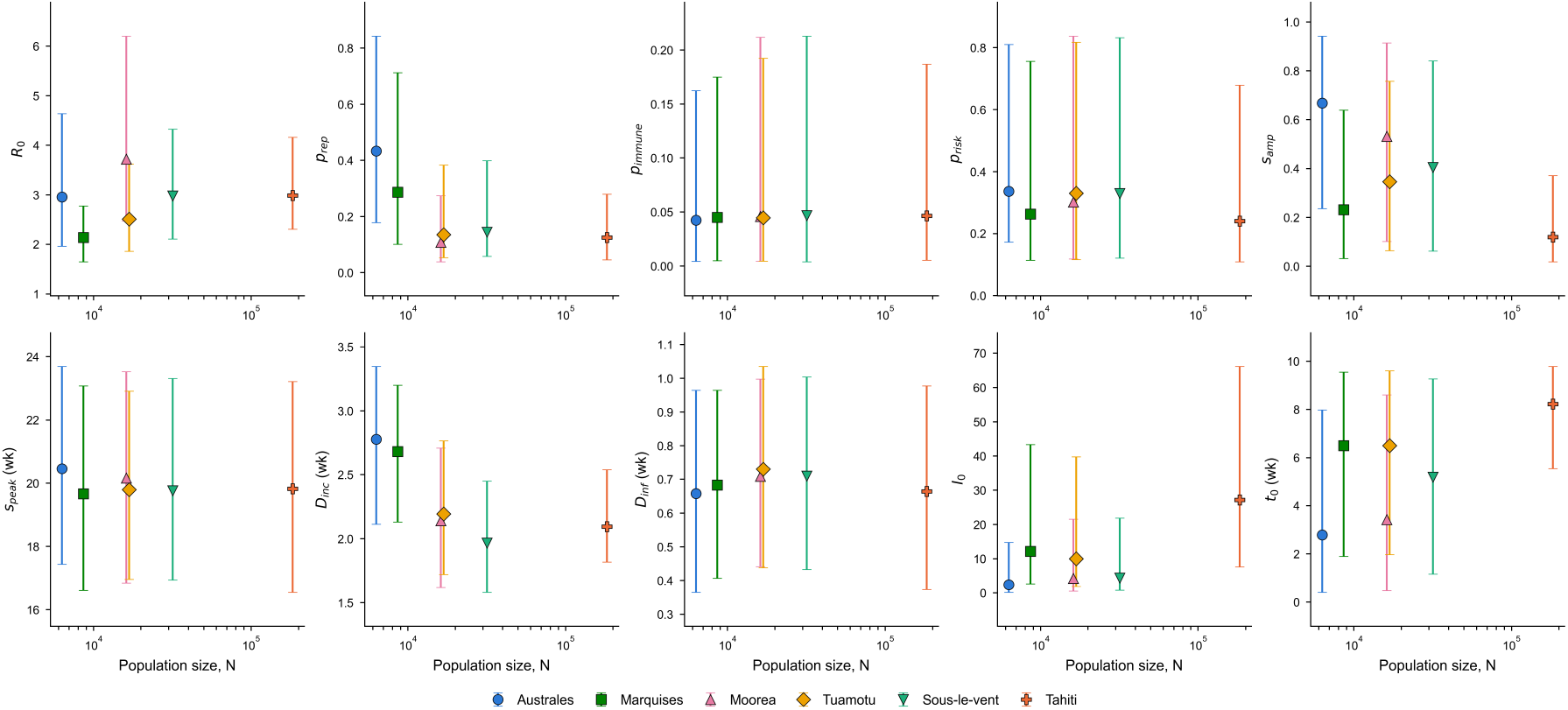
Posterior estimates of model parameters as a function of population size, shown for each of the six French Polynesia archipelagos. Points indicate posterior medians; error bars show 5th and 95th percentiles of the marginal posterior distribution.

**SI Fig. 2:**
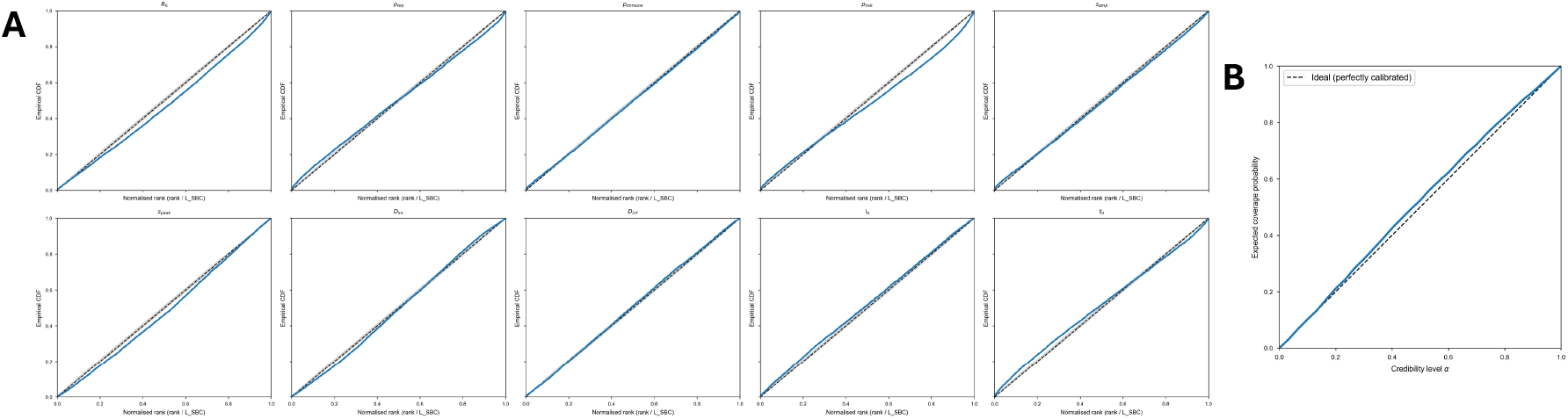
(A) Cumulative distribution of ranks for individual parameters for Zika transmission model. For each simulation, the rank is calculated using 1000 samples from the estimated posterior. (B) Expected coverage probability vs credibility level as calculated from the TARP analysis.

**SI Fig. 3:**
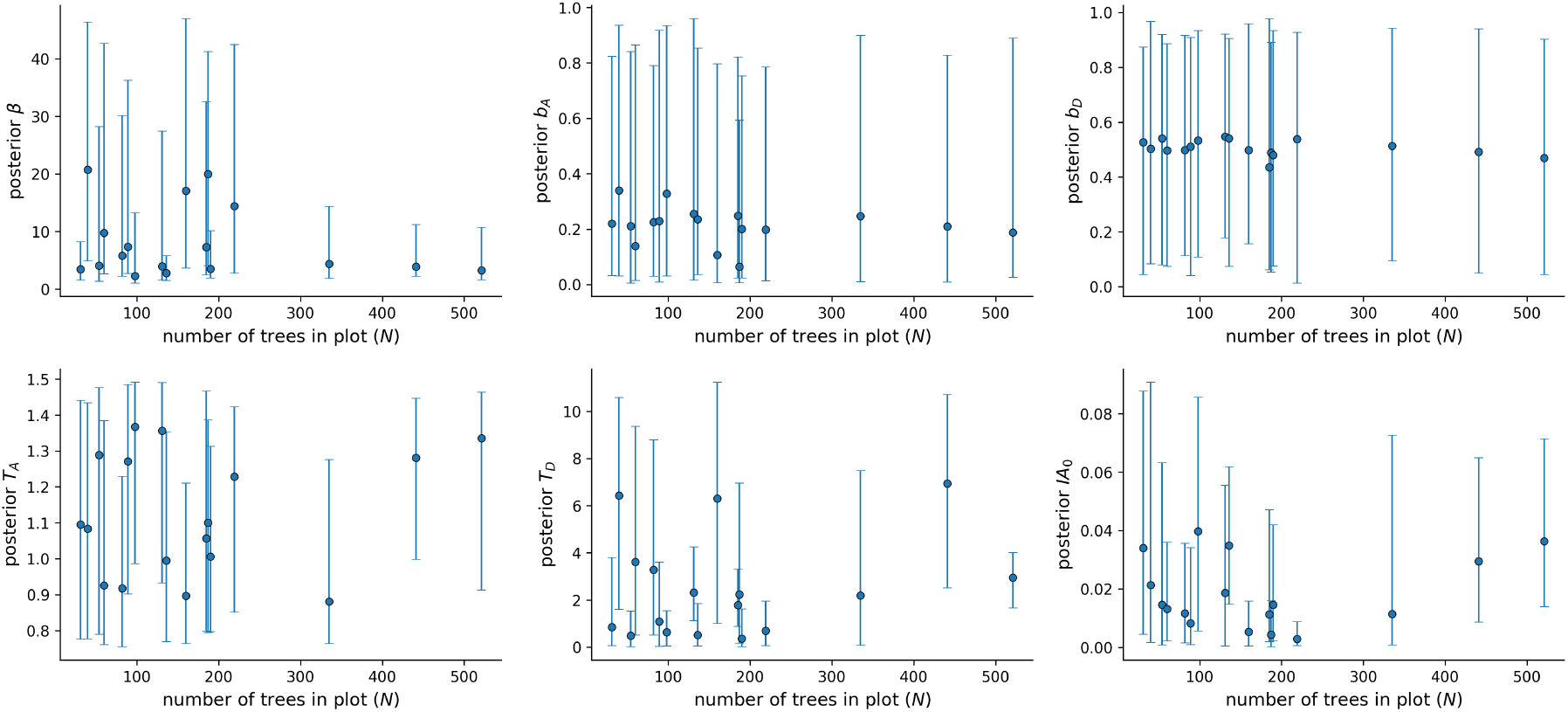
Posterior estimates of the *X. fastidiosa* transmission model parameters as a function of population size, shown for each of the groves. Points indicate posterior medians; error bars show 5th and 95th percentiles of the marginal posterior distribution.

**SI Fig. 4:**
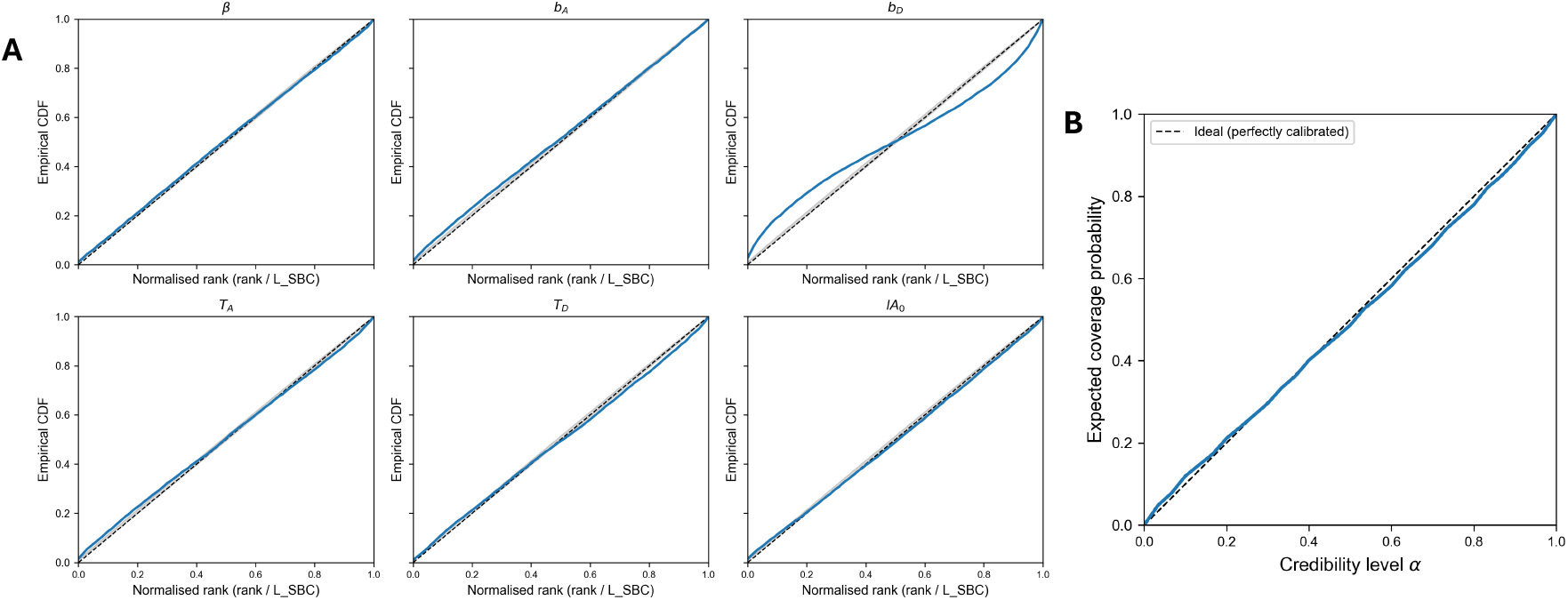
(A) Cumulative distribution of ranks for individual parameters for *xf* transmission model. For each simulation, the rank is calculated using 1000 samples from the estimated posterior. (B) Expected coverage probability vs credibility level as calculated from the TARP analysis.

**Table SI1.** Prior distribution ranges for Zika model parameters.

| Parameter | Description | Min | Max |
| --- | --- | --- | --- |
| $R_0$ | Basic reproduction number | 0.00 | 25.00 |
| $p_{\text{rep}}$ | Proportion of cases reported | 0.00 | 1.00 |
| $p_{\text{immune}}$ | Proportion immune | 0.00 | 0.35 |
| $p_{\text{risk}}$ | Proportion at risk | 0.10 | 1.00 |
| $s_{\text{amp}}$ | Amplitude of seasonality | 0.00 | 1.00 |
| $s_{\text{peak}}$ | Peak timing of seasonality | 1.00 | 30.00 |
| $D_{\text{inc}}$ | Intrinsic incubation period (weeks) | 1.50 | 3.60 |
| $D_{\text{inf}}$ | Infectious period (weeks) | 0.20 | 1.25 |
| $I_0$ | Initial infectious humans | 0.00 | 100.00 |
| $t_0$ | Initial time offset | 0.00 | 10.00 |

**Table SI2.** Prior distribution ranges for the *Xylella fastidiosa* epidemiological model parameters.

| Parameter | Description | Min | Max |
| --- | --- | --- | --- |
| $\beta$ | Contact rate of symptomatic trees (per year) | 0.00 | 50.00 |
| $b_A$ | Relative infectivity of asymptomatic trees | 0.00 | 1.00 |
| $b_D$ | Relative infectivity of desiccated trees | 0.00 | 1.00 |
| $T_A$ | Mean symptomless period (years) | 0.75 | 1.50 |
| $T_D$ | Mean time to desiccation (years) | 0.00 | 12.00 |
| $I_{A,0}$ | Initial proportion infected without symptoms | 0.00 | 0.10 |

